# Single-molecule analysis of purified proteins and nuclear extracts: insights from 8-oxoguanine glycosylase 1

**DOI:** 10.1101/2023.11.01.565178

**Authors:** Matthew A. Schaich, Tyler M. Weaver, Vera Roginskaya, Bret D. Freudenthal, Bennett Van Houten

## Abstract

By observing one molecule at a time, single-molecule studies can offer detailed insights about biomolecular processes including on rates, off rates, and diffusivity of molecules on strands of DNA. A recent technological advance (Single-molecule Analysis of DNA-binding proteins from Nuclear Extracts, SMADNE) has lowered the barrier to entry for single-molecule studies, and single-molecule dynamics can now be determined directly out of nuclear extracts, providing information in an intermediate environment between purified proteins in isolation and the heterogeneity of a nucleus. To compare and contrast the single-molecule DNA binding dynamics in nuclear extracts versus purified proteins, combined optical tweezers and fluorescence microscopy experiments were performed with purified GFP-tagged 8-oxoguanine glycosylase 1 (OGG1), purified GFP-OGG1 spiked into nuclear extracts, and nuclear extracts from human cells overexpressing GFP-OGG1. We observed differences in undamaged DNA binding during DNA damage search in each of the three conditions. Purified GFP-OGG1 engaged undamaged DNA for a weighted average lifetime of 5.7 s and 21% of these events underwent DNA diffusion after binding. However, unlike other glycosylases studied by SMADNE, OGG1 does not bind non-damaged DNA efficiently in nuclear extracts. In contrast, GFP-OGG1 binding dynamics on DNA substrates containing oxidative damage were relatively similar in all three conditions, with the weighted average binding lifetimes varying from 2.2 s in nuclear extracts to 7.8 s with purified GFP-OGG1 in isolation. Finally, we compared the purified protein and nuclear extract approaches for a catalytically dead OGG1 variant (GFP-OGG1-K249Q). This variant greatly increased the binding lifetime for oxidative DNA damage, with the weighted average lifetime for GFP-OGG1-249Q in nuclear extracts at 15.4 s vs 10.7 s for the purified protein. SMADNE will provide a new window of observation into the behavior of nucleic acid binding proteins only accessible by biophysicists trained in protein purification and protein labeling.

## Introduction

When proteins are properly purified, experimental results hold the distinct advantage of directly observing protein behavior without concern that unknown factors influence the results. Furthermore, protein purification has previously been an obligate requirement for numerous types of biophysical analyses, ranging from enzyme kinetics and structural studies to experiments where protein behavior is monitored at the single-molecule level [1–4]. With the advent of new single-molecule techniques [5–12], such as the Single-molecule Analysis of DNA-binding proteins from Nuclear Extracts (SMADNE), the necessity to always purify proteins has been lifted in order to study proteins at the single-molecule level. Utilizing nuclear extracts directly expressed from mammalian cells has a number of advantages such as: rapidly screening variants, post-translational modifications (PTMs) can be preserved, proteins that are traditionally challenging [13, 14] to express/purify from bacteria can be readily obtained within minutes of lysing cells, purification of large mammalian complexes is readily amenable from extracts, and using lysates does not require the hours if not days of time necessary to fully purify protein. At the same time, experiments like SMADNE that use nuclear extracts hold the dual advantage and disadvantage that many of the thousands of proteins present in a nucleus are also present in the experiment, albeit at ∼500-fold lower concentrations than in the nucleus, which may make the results difficult to interpret. Though, the presence of other proteins is also beneficial because the experimental results may be more indicative of behavior in a biological context compared to a protein studied in isolation. However, this reward holds the caveat that the identities of these unseen “dark” proteins are unknown until follow-up experiments are performed, such as fluorescent labeling or knocking down putative interacting partners [5].

8-oxoguanine glycosylase 1 (OGG1) is a key protein involved in the repair of the oxidative damage 8-oxoguanine (8-oxoG) during base excision repair (BER), where OGG1 identifies 8-oxoG across from a cytidine and subsequently cleaves the glycosidic bond to leave behind an abasic site [15]. Similar to all DNA glycosylases, OGG1 faces the extraordinary challenge of rapidly identifying 8-oxoG amongst billions of undamaged DNA base pairs [16]. Thus, it has been proposed and observed that OGG1 diffuses along the DNA helix to aid in its search for damage [17, 18]. The most direct way to understand the damage search process of OGG1 is fluorescent labeling of the protein and observing its movement on DNA in real time. Importantly, OGG1 (and bacterial DNA-formamidopyrimidine glycosylase, FPG) have been extensively characterized at the single-molecule level using multiple different imaging techniques [5, 17, 19]. Additionally, OGG1 has also been characterized on a variety of nucleic acid substrates including undamaged DNA, DNA containing abasic sites [18], and DNA containing oxidative damage [5]. Finally, OGG1 has been labeled with numerous fluorescent labeling strategies, including Cy3 maleimide labeling, Qdot conjugation with an antibody, and fusing a fluorescent fusion protein to the protein of interest, making it a great model system for single-molecule studies [5, 17, 18].

The purpose of this article is two-fold, to review the SMADNE approach and to compare the behavior of a purified protein versus the SMADNE approach using OGG1. In our experiments, we utilized OGG1-GFP as a model system to determine how nuclear proteins present in extracts may alter single-molecule binding kinetics. Three different contexts were assayed: (1) OGG1-GFP purified from a bacterial expression system, (2) purified OGG1-GFP spiked into the nuclear extracts with no overexpressed protein, and (3) OGG1-GFP overexpressed in nuclear extracts without purification (SMADNE approach). This has allowed us to directly compare and contrast these three different experimental systems, while also beginning to deconvolute the role of unknown nuclear proteins in the DNA damage search and detection process.

## Materials and methods

### Purification of DNA Glycosylases

Human OGG1 WT or OGG1 K249Q in a pET-His6-GFP-TEV bacterial expression vector (Addgene #29663) were obtained from GenScript. All GFP-OGG1 proteins were expressed in BL21-CodonPlus (DE3)-RP *Escherichia coli* (*E. coli*) competent cells (Agilent). The cells were grown at 37 °C to an OD600-0.6 and OGG1 expression was induced with 0.5 mM isopropyl-b-D-thiogalactopyranoside (IPTG) for 18 hours at 18 °C. The cells were harvested and lysed via sonication in a buffer containing 150mM NaCl, 50mM HEPES (pH-7.5), and an Inhibitor Cocktail (Benzamidine, Leupeptin, AEBSF, Pepstatin A). The cell lysate was cleared for 1□h at 24,242□×□g. The resulting supernatant containing GFP-OGG1 protein was purified via two 5 mL HisTrap HP (Cytiva) equilibrated with 150mM NaCl, 50mM HEPES (pH-7.5), and 20 mM Imidazole, and the protein eluted using a linear gradient with a buffer containing 150mM NaCl, 50mM HEPES (pH-7.5), and 500 mM Imidazole. The GFP-OGG1 proteins was further purified via cation-exchange chromatography using a RESOURCE S column (Cytiva) equilibrated in 50mM NaCl, 50mM HEPES (pH-7.5), 1 mM DTT, and 1 mM EDTA, and the protein eluted off the column using a linear gradient to a final buffer containing 1 M NaCl, 50mM HEPES (pH-7.5), 1 mM DTT, and 1 mM EDTA. The GFP-OGG1 protein was then polished via size exclusion chromatography using a HiPrep Sephacryl S-200 HR (16/60) in a buffer containing 150mM NaCl, 50mM HEPES (pH-7.5), and 1 mM TCEP. The purified GFP-OGG1 proteins were concentrated to ∼100 μM and stored long term at −80 °C

The untagged OGG1 WT and K249Q proteins were generated from the GFP-OGG1 WT and K249Q proteins. In brief, the untagged OGG1 protein was liberated from the GFP-tag using Tobacco Etch Virus (TEV) protease (1 mg TEV per 10mg OGG1) in a buffer containing 50mM NaCl, 50mM HEPES (pH-7.5), and 1 mM DTT. The resulting untagged OGG1 proteins were subsequently purified by cation-exchange chromatography using a RESOURCE S column (Cytiva) equilibrated in 50mM NaCl, 50mM HEPES (pH-7.5), 1 mM DTT, and 1 mM EDTA, and the protein eluted off the column using a linear gradient to a final buffer containing 1 M NaCl, 50mM HEPES (pH-7.5), 1 mM DTT, and 1 mM EDTA. The untagged OGG1 protein was further purified via size exclusion chromatography using a HiPrep Sephacryl S-200 HR (16/60) in a buffer containing 150mM NaCl, 50mM HEPES (pH-7.5), and 1 mM TCEP. The purified untagged OGG1 proteins were concentrated to ∼100 μM and stored long term at −80 °C

### DNA Glycosylase Activity Assays

For DNA glycosylase activity assays, a double stranded 8-oxoG DNA substrate was generated from the following oligonucleotides: 5′-/FAM/CTG-CAG-CTG-ATG-CGC-C(8-oxo-dG)T-ACG-GAT-CCC-CGG-GTAC-3′ and 5′-GTA-CCC-GGG-GAT-CCG-TAC-GGC-GCA-TCA-GCT-GCAG-3′ (undamaged strand). The oligonucleotides were annealed in a buffer containing 10 mM Tris (pH-8.0) and 1 mM EDTA by heating to 90 °C for 5 min, and subsequently cooling to 4 °C using a rate of −1 °C min^−1^.

DNA glycosylase assays were carried out by incubating each respective purified OGG1 protein (20 nM) with 8-oxoG DNA (20 nM) in a buffer containing 100 mM KCl, 50 mM HEPES (pH-7.5), 100 μg/mL BSA, and 1 mM DTT. The reactions were quenched after 15 mins through the addition of a 1 M NaOH, boiled at 95°C for 5 minutes to cleave the OGG1 product AP site, and the reactions neutralized with 1 M HCl. The reactions were subsequently mixed with a 1:1 v/v ratio of formamide loading dye (100 mM EDTA, 80% deionized formamide, 0.25 mg/ml bromophenol blue, 0.25 mg/ml xylene cyanol, and 8 M urea), and resolved on a 15% denaturing polyacrylamide gel. The resolved substrate (8-oxoG) and product (AP site) bands were then visualized using an Amersham Typhoon RGB imager.

### Cell lines

Dulbecco’s Modified Eagle Medium (DMEM) supplemented with 4.5g/l glucose, 10% fetal bovine serum (Gibco), 5% penicillin/streptavidin (Life Technologies) was utilized to culture U2OS cells in conditions with 20% oxygen. Transfection and nuclear extraction were performed as in the SMADNE methodology [5]. Briefly, four µg of plasmid per four million cells as a transfection with lipofectamine 2000. To prepare the nuclear extract control samples, the same lipofectamine protocol was followed but no plasmid was added. After a period of 24h nuclear extracts (∼80 μL) were generated using a kit from Abcam (ab113474). Resultant nuclear extracts were aliquoted into single-use tubes (∼10 μL each at ∼1 mg/mL) and flash frozen in liquid nitrogen prior to storing them at −80 °C. The frozen extracts are stable for at least 12 months.

### DNA substrate generation

Lambda DNA for C-trap experiments was purchased from New England Biolabs and its overhangs were biotinylated with biotinylated dCTP as performed previously [5]. Oxidative damage was introduced by incubating with 0.2 μg/mL methylene blue (as performed here [19]) and exposed to 660 nm light for 10 minutes. Based on the previous work, we estimate this protocol introduces 1 damaged base per ∼440 bp throughout the length of the lambda DNA.

### Single-molecule experiments

**Equipment:** All single-molecule experiments were performed on a LUMCKS C-trap ® [5] instrument, an optical platform that combines optical tweezers, a three confocal fluorescence microcopy, and a five chamber microfluidic flow cell [5]. Excitation lasers were used at 5% and emission was monitored using single photon detectors during kymograph acquisition at 10 frames per second and 100 nm pixels in the Y-axis.

### DNA tether formation and positioning

Utilizing four channels of the microfluidic flow cell, experimental design consisted of four major steps prior to imaging. First, after opening the valves of the flow cell and pressurizing to 0.3 bar to maintain laminar flow, streptavidin-coated polystyrene beads (4.4-4.8 micron) were immobilized in two separate optical traps. Then the beads were moved to the second channel of the flow cell where the biotinylated lambda DNA was flowing. By varying the distance between the beads between 10 microns to 15 microns and monitoring the force compared to an extensible worm-like chain model, we confirmed that a single DNA tether was obtained between the two beads. Then the tethered DNA was moved to a channel containing buffer that consisted of 150 mM NaCl, 20 mM HEPES pH 7.5, 5% glycerol, 0.1 mg/mL BSA, 1 mM freshly thawed DTT, and 1 mM Trolox. The DNA was washed for ten seconds before moving to the channel with the fluorescent GFP-OGG1 (either as purified proteins at 20 nM concentration, 10 nM purified protein spiked into nuclear extracts without overexpression diluted 1:10 in imaging buffer, or nuclear extracts diluted 1:10 in imaging buffer to ∼ 0.1 mg/ml), pulling the tension to 10 pN, and collecting binding events along the DNA. For the experiments containing nuclear extracts, buffer and nuclear extracts were flowed in fresh every five minutes. For experiments with purified proteins, the sample was refreshed more frequently to account for the decay in fluorescent intensity, typically every 1-2 minutes and when binding events were no longer occurring.

### Confocal imaging

GFP signals were collected by exciting with a 488 nm laser at 5% power (∼2 µW at the objective) and emission was collected through a 500-550 nm band pass filter. Imaging was performed with a 1.2 NA 60X water objective and intensities measured with single-photon avalanche photodiode detectors. Kymograph scans were collected along the length of the DNA and 10 frames per second with a pixel size of 100 nm and exposure time of 0.1 msec per pixel. In the case of OGG1-249Q-GFP on undamaged DNA, this time resolution made line tracking difficult given the short binding lifetime, so the frame rate was increased to 33 frames per second.

### Data analysis

Kymographs were analyzed with custom software from Lumicks (Pylake). Images for publication were generated with the .h5 Visualization GUI (2020) by John Watters, accessed through harbor.lumicks.com. As GFP has been previously observed to blink up to two seconds[20], any events occurred at the same position with less than two seconds of non-fluorescent time between them were connected and counted as a single binding event.

Motile events were analyzed by extracting the mean square displacement utility of Pylake, where the plots for each lag time were exported for custom fitting. The equation utilized is shown below:

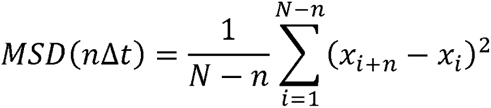

where *N* is total number of frames in the phase, *n* is the number of frames at a given time step, Δ*t* is the time increment of one frame, and *x_i_* is the particle position in the *i*^th^ frame. The diffusion coefficient (*D*) was determined by fitting a model of one-dimensional diffusion to the linear portion of the MSD plots:

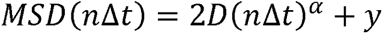

where α is the anomalous diffusion coefficient and *y* is a constant (*y*-intercept). Each plot was analyzed using Graphpad Prism, and the maximum time window adjusted to include as much of the linear portion of the graph as possible. Fittings resulting in *R^2^* less than 0.8 or using less than 10% of the MSD plot were excluded.

## Results

### Purified OGG1 scans undamaged DNA for damage

To establish the search mechanisms of OGG1-GFP on DNA substrates with defined tension, we expressed and purified a GFP-tagged OGG1 from bacterial overexpression (Fig. S1 and 1A). Importantly, the activity of OGG1 in the presence and absence of the GFP tag are virtual identical, indicating the GFP-label does not interfere with the ability of OGG1 to excise an 8-oxoG:C base pair in DNA (Fig. S1). To establish DNA tethers for visualization, we utilized a LUMICKS C-trap®, which combines a five chamber flow cell with optical tweezers and a fluorescence microscope (Figure 1B) [5]. This system enables us to suspend 48.5 kb of dsDNA in a flow chamber with precise force measurement and control. The DNA tethering is initially performed in an isolated channel prior to moving the DNA tether into a new channel containing the protein of interest to study its behavior. The DNA was positioned in the middle of the flow channel away from the surface of the glass, which has proven crucial for preventing imaging difficulties caused by debris nonspecifically adhering to the glass of the flow cell.

**Fig. 1:**
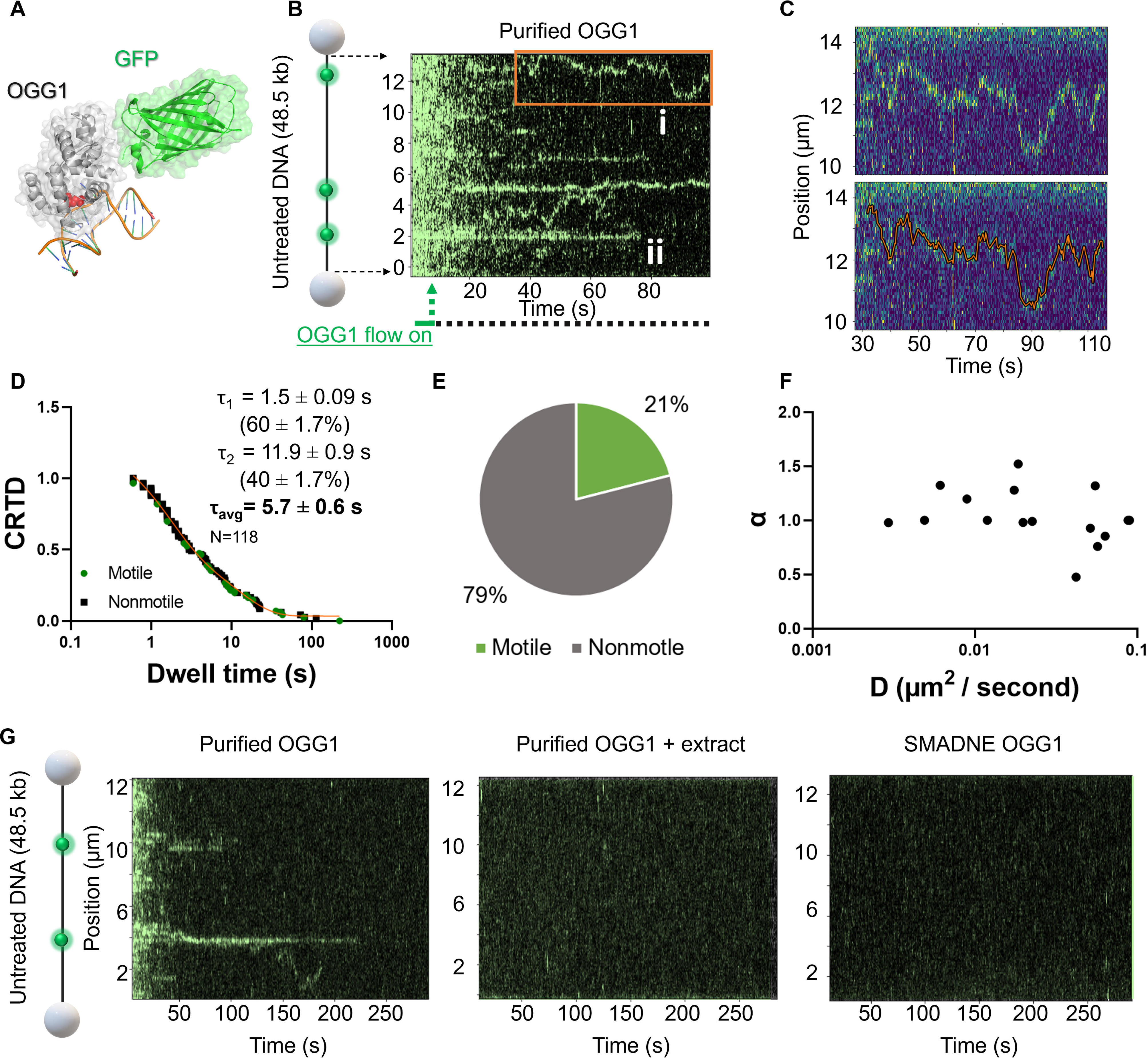
OGG1-GFP binds undamaged DNA with multiple modes. (A) A structural model of GFP-tagged OGG1 bound to damaged DNA, from PDB codes 1YQR and 5LK4. (B) A representative kymograph of OGG1-GFP binding undamaged DNA, with a cartoon on the left showing the positions of the beads and DNA. The times at which microfluidic flow was present are also indicated. (C) A representative motile OGG1-GFP event with line tracking shown beneath the raw kymograph. (D) A cumulative residence time distribution (CRTD) is shown. The fit of double-exponential decay functions is shown in orange, and nonmotile dwell times shown in black, and motile dwell times shown in green. (E) The distribution of motile to nonmotile events. (F) Diffusivity and anomalous diffusion coefficient for motile OGG1-GFP events. (G) Representative five-minute kymographs for purified OGG1-GFP, purified OGG1-GFP plus nontransfected nuclear extract, and OGG1-GFP generated in mammalian cells prior to nuclear extraction.

Upon moving the tethered DNA into the channel of the flow cell containing purified OGG1-GFP, we observed a variety of single-molecule binding events across the length of the DNA. This includes binding events where OGG1-GFP engages the DNA and remained static and binding events where OGG1-GFP appeared to diffuse on the DNA prior to dissociation (Fig. 1B, 1C, Table 1). When presented as a kymograph (with each pixel in the x-axis representing 100 ms and each pixel in the y axis representing 100 nm), stationary binding events and localized searching events (beneath the localization precision of the instrument) appear as straight green lines on the DNA, whereas long-range motile events appear as jagged lines as OGG1-GFP diffuses along the DNA over time. Surprisingly, we observed a rapid reduction in the background fluorescence within 15 −20 seconds generated from OGG1-GFP molecules diffusing in solution and not bound to the DNA. This wave of fluorescence reduced relatively quickly after flowing in fresh protein – as the valves were sealed shut to the flow cell. This reduction in the available protein (fading phenomenon) potentially caused by molecules sticking to the glass outside of the imaging plane, reducing the total amount of protein available for binding – however this is likely protein and tag dependent, as previous reports with other purified proteins using other labels did not observe this phenomenon [21, 22]. In addition, this fading phenomenon resulted in the majority of binding events occurring within the first few seconds of a kymograph.

**Table 1:** Summary of single-molecule binding kinetics in this study.

| DNA tether type | OGG1 variant | Purified protein lifetime (s) | Purified + extract lifetime (s) | SMADNE lifetime (s) |
| --- | --- | --- | --- | --- |
| Undamaged | WT | $\tau_1 = 1.5 \pm 0.09$ s<br>$(60 \pm 1.7\%)$<br>$\tau_2 = 11.9 \pm 0.9$ s<br>$(40 \pm 1.7\%)$<br>$\tau_{avg} = 5.7 \pm 0.6$ s * | No events | No events |
| | K249Q | $\tau = 0.47 \pm 0.14$ s | No events | No events |
| 8-oxoguanine | WT | $\tau_1 = 4.4 \pm 0.9$ s<br>$(46 \pm 18\%)$<br>$\tau_2 = 10.6 \pm 2.2$ s<br>$(54 \pm 18\%)$<br>$\tau_{avg} = 7.8 \pm 2.2$ s | $\tau_1 = 2.0 \pm 0.05$ s<br>$(88 \pm 1.8\%)$<br>$\tau_2 = 45.1 \pm 22$ s<br>$(12 \pm 1.8\%)$<br>$\tau_{avg} = 7.1 \pm 3.5$ s | $\tau_1 = 0.8 \pm 0.06$ s<br>$(51\% \pm 3.9)$<br>$\tau_2 = 3.2 \pm 0.25$ s<br>$(49 \pm 3.9\%)$<br>$\tau_{avg} = 2.0 \pm 0.2$ s |
| | K249Q | $\tau_1 = 4.7 \pm 0.9$ s<br>$(46 \pm 11\%)$<br>$\tau_2 = 15.8 \pm 2.6$ s<br>$(54 \pm 11\%)$<br>$\tau_{avg} = 10.7 \pm 2.7$ s | $\tau_1 = 2.9 \pm 0.3$ s<br>$(52 \pm 2.7\%)$<br>$\tau_2 = 24.8 \pm 4.6$ s<br>$(48 \pm 2.7\%)$<br>$\tau_{avg} = 13.4 \pm 2.9$ s | $\tau_1 = 7.7 \pm 0.3$ s<br>$(78 \pm 2.3\%)$<br>$\tau_2 = 42.9 \pm 8.7$ s<br>$(22 \pm 2.3\%)$<br>$\tau_{avg} = 15.4 \pm 3.2$ s |

\* = 21% motile events

Upon tracking the duration of binding events, we observed that dwell times occurred over a wide range, from transient events that occurred in less than one second to long-lived events that lasted over 100 seconds (Fig. 1D). These events were sorted by duration and fit to a cumulative residence time distribution (CRTD) plot [5]. Upon fitting to a double-exponential decay function, the events exhibited two binding lifetimes, one at 1.5 s (60% contributing) and one at 11.9 s (40% contributing). These two different binding lifetimes could be a result of conformational proofreading by OGG1, where one protein state acts as a brief DNA sampling conformation and the second conformation resides longer on the DNA [23]. Alternatively, the fast phase could be non-specific binding, and the longer-lived binding events may represent cryptic lesions that were introduced into the lambda DNA during purification and processing prior to stringing up in the C-trap. Of the binding events observed, 21 % exhibited motile behavior (Fig. 1E), diffusing on the DNA before dissociating. The diffusivity of the motile events was determined using mean square displacement (MSD) analysis and was on average 0.035 μm^2^/s (Fig. 1F). This average diffusivity value was much slower than the 0.58 μm^2^/s previously reported for Cy3 labeled OGG1, which could be explained in part by the 100 μm/s flow velocity of the previous collection, compared to our data collection without flow [17]. In contrast to the binding events observed with purified OGG1-GFP on undamaged DNA, we did not observe binding events when the purified OGG1-GFP was spiked into nuclear extract or when using the SMADNE approach, although the background fluorescence intensity was more consistent than with purified protein alone (Fig. 1G). Thus, we could not detect any 1D diffusion by OGG1-GFP on the undamaged DNA in the presence of nuclear extracts, suggesting nuclear proteins can alter the OGG1 search mechanism on DNA.

### OGG1 robustly binds 8-oxoG as a purified protein and in the presence of nuclear extract

To assess the ability of OGG1-GFP to bind 8-oxoG, we exposed the lambda DNA to methylene blue and 660 nm light to generate oxidative damage, primarily 8-oxoG, throughout the length of the DNA sequence (approximately every 440 bp) as performed previously [19]. With this damage load, we no longer observed motile binding events with purified OGG1-GFP. This could be due to higher affinities for 8-oxoG over non-damaged DNA so that 3D diffusion is sufficient for a binding event, or OGG1 does not need to scan very far before encountering a damage site, since 440 bp is only 1.3 pixels in the kymograph obtained from the C-trap (Fig. 2A). All three conditions exhibited a wide range of dwell times, and the purified OGG1-GFP bound with a lifetimes of 4.4 s (46%) and 10.6 s (54%), for a weighted average lifetime of 7.8 s. When the purified OGG1-GFP was spiked into nuclear extract, we observed binding events with two dwell times corresponding to 2.0 s (88%) and 45.1 s (12%), similar to the behavior with purified OGG1-GFP (Fig. 2B). With this condition, the slow off rate was both slower and less frequent than the purified protein alone, which may implicate that extract proteins stabilize OGG1-GFP events that engage DNA damage. Interestingly, we did not observe the previously described fading phenomenon seen with purified OGG1-GFP, suggesting the presence of other proteins may help to prevent OGG1-GFP from sticking to the glass of the flow cell or that chaperones present may help stabilize the protein. Finally, we tested OGG1-GFP from the SMADNE approach, and found that it also exhibited exclusively nonmotile events on the damaged DNA. While the range of dwell times were less than a second to over 100 seconds, many short binding events cause the CRTD plot to exhibit two shorter lifetimes, one at 0.8 s (51%) and one at 3.2 s (49%), for a weighted average lifetime of 2.0 s (Fig. 2C). These relatively short dwell times for OGG1-GFP prepared in human cell nuclear extracts suggests that OGG1-GFP in extracts is altered in some way to change its off rate, which could be caused by post-translational modifications, or downstream enzymatic processing of the OGG1-DNA complex (i.e., APE1).

**Fig. 2:**
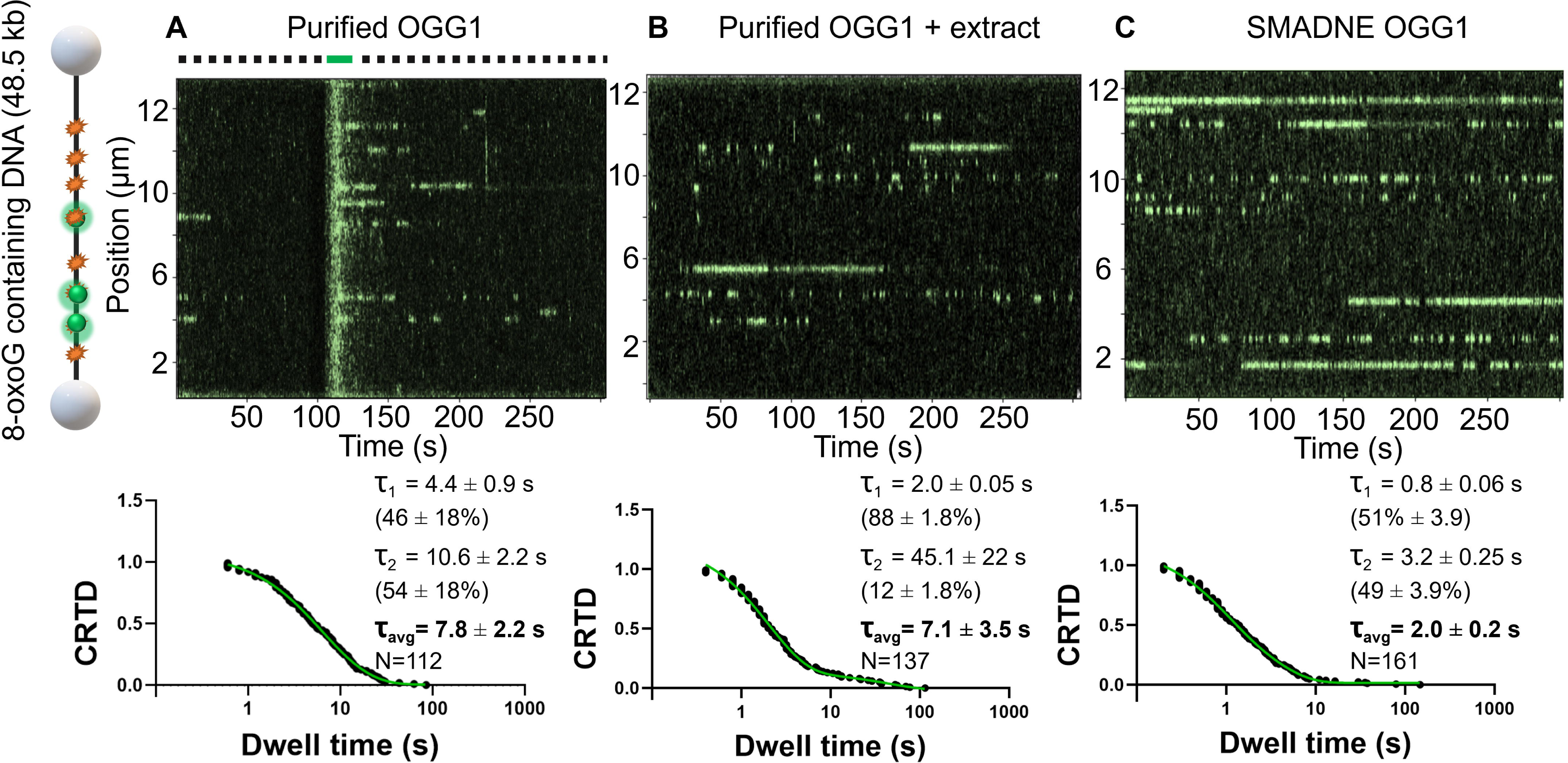
The impact of proteins in nuclear extracts on OGG1 binding damaged DNA. (A) A representative kymograph of purified OGG1-GFP binding DNA treated with methylene blue and light to form 8-oxoguanine. Schematic on left shows positions of the beads and DNA. CRTD plot for purified OGG1 on damaged DNA is also shown. (B) A representative kymograph of purified OGG1-GFP spiked into nuclear extracts is shown in green, and the resultant CRTD plot is displayed below. (C) Kymograph obtained with the single-molecule analysis of DNA-binding proteins from nuclear extracts (SMADNE) approach with OGG1-GFP overexpressed, with the resultant CRTD plots and fits shown underneath.

### Catalytically dead OGG1 transiently engages undamaged DNA

To better understand how nuclear extracts impact binding lifetimes of proteins and determine the utility of each technique in studying protein variants, we performed additional experiments with a catalytically dead OGG1 variant (K249Q). This variant replaces the positively charged lysine, which serves as a nucleophile during 8-oxoG glycosidic bond cleavage, with a glutamine residue (K249Q). This renders the OGG1 K249Q mutant enzymatically inactive, though it maintains the ability to robustly engage 8-oxoG [24]. On undamaged DNA, binding events were evident with purified OGG1-K249Q-GFP, but we observed a much shorter binding lifetime than WT OGG1-GFP, fitting to a single-exponential decay function with a lifetime of 0.47 s. Furthermore, no visibly motile events were observed with this catalytic mutant. Thus, residue K249Q plays a role in productive search mechanisms by OGG1 as well as its role in catalysis. Potentially the impaired searching of OGG1 could be explained by decreased engagement on the DNA: if the sampling events on the DNA are too transient to establish long range motility on the DNA, then the effectiveness of a search will be reduced. Although binding events were observed for the undamaged DNA with purified OGG1-K249Q-GFP (Fig. 3A-B), events on undamaged DNA were not observed when the purified OGG1-K249Q-GFP was spiked into nuclear extract or expressed via the SMADNE approach (Fig. 3C-D).

**Fig. 3:**
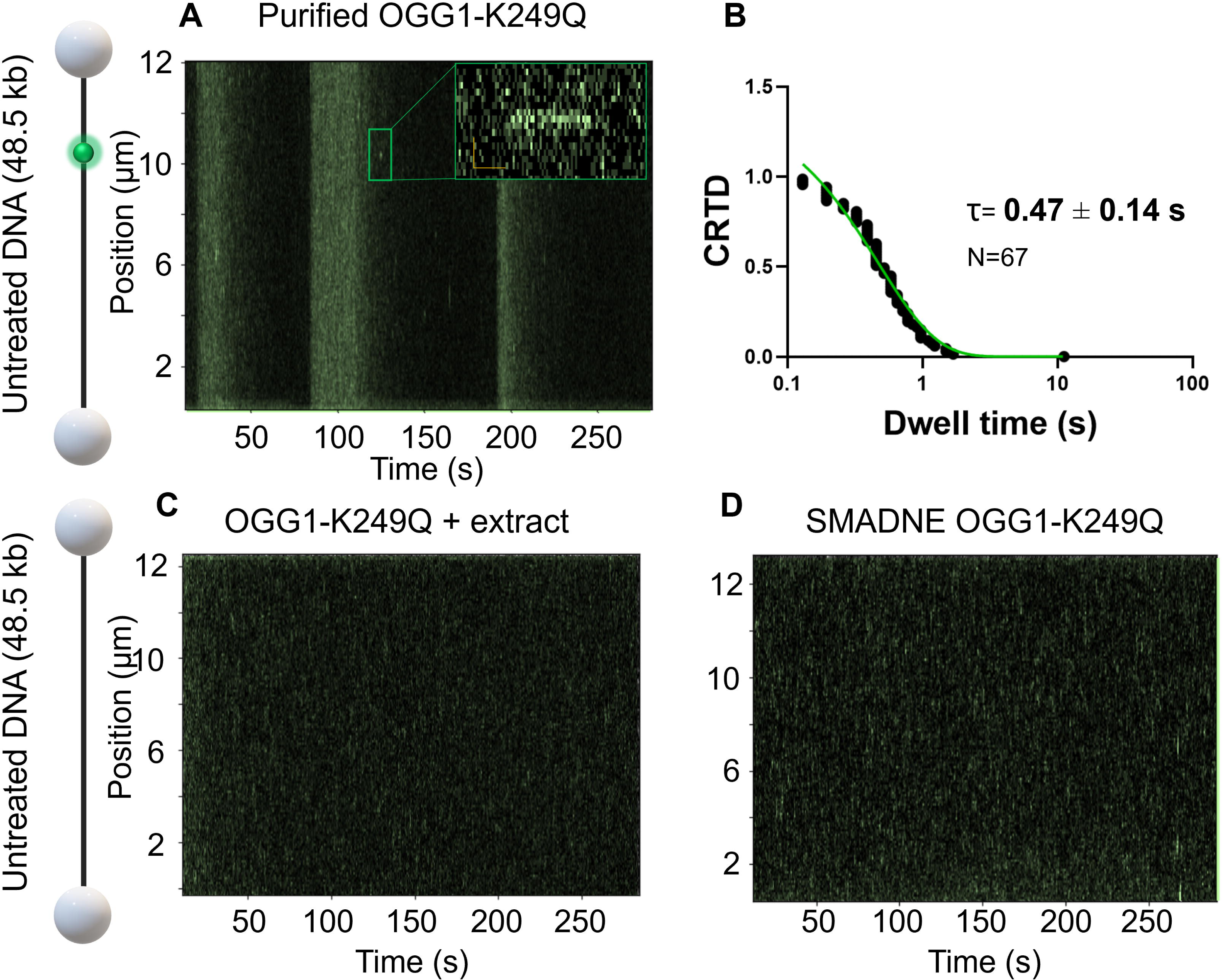
Catalytically dead OGG1 engages undamaged DNA. (A) Undamaged DNA was incubated with purified OGG1-K249Q-GFP, and transient interactions were observed (shown in green). (B) CRTD plot from the dwell times observed is displayed with a single-exponential decay fit. (C) On a similar timescale, events were not observed when the purified protein was spiked into nuclear extracts or (D) when the sample was generated with SMADNE.

### OGG1-K249Q-GFP engages damage sites with longer lifetimes than WT OGG1

The unique properties of the OGG1 K249Q variant also allows us to unambiguously determine binding dynamics during 8-oxoG recognition, rather than a combination of binding events associated with 8-oxoG and abasic sites that are generated following OGG1 catalysis. When we tested the binding behavior of catalytically-dead OGG1-K249Q-GFP on damaged DNA, we observed long-lived binding events in all three experimental conditions (i.e., purified protein, purified protein plus nuclear extract, and SMADNE, Fig 4). This trend recapitulates the behavior that we previously observed with WT OGG1-GFP, where the presence of nuclear extracts reduced non-specific binding events but still allowed for successful engagement of DNA damage. In the case of purified OGG1-K249Q-GFP, we observed exclusively nonmotile events for this substrate, similar to what was observed for WT OGG1-GFP on DNA containing 8-oxoG (Fig. 4A). These events exhibited dwell times that fit to a double-exponential decay function, with one lifetime at 4.7 s (46%) and the other at 15.8 s (54%), for a weighted average lifetime of 10.7 s. Thus, there was a 20-fold increase in the binding lifetime of OGG1-K249Q-GFP between undamaged DNA and DNA containing 8-oxoG. For the purified OGG1-K249Q-GFP spiked into nuclear extract, similar binding lifetime and behavior was observed, with exclusively nonmotile events. Dwell times fit a double-exponential decay function with one lifetime at 2.9 s and one lifetime at 24.8 s, with the short lifetime contributing 52% and a weighted average lifetime of 13.4 s (Fig. 4B). Finally, binding events from the OGG1-K249Q-GFP SMADNE approach exhibited two dwell times, with one lifetime at 7.7 s and the second at 42.9 s, with the fast lifetime contributing 78% (Fig. 4C). These dwell times yield a weighted average lifetime for the SMADNE OGG-K249Q-GFP of 15.4 s, which is similar to the lifetimes of the other two conditions. In summary, we observed a robust increase in dwell times for OGG-K249Q-GFP binding 8-oxoG in all three experimental regimes.

**Fig. 4:**
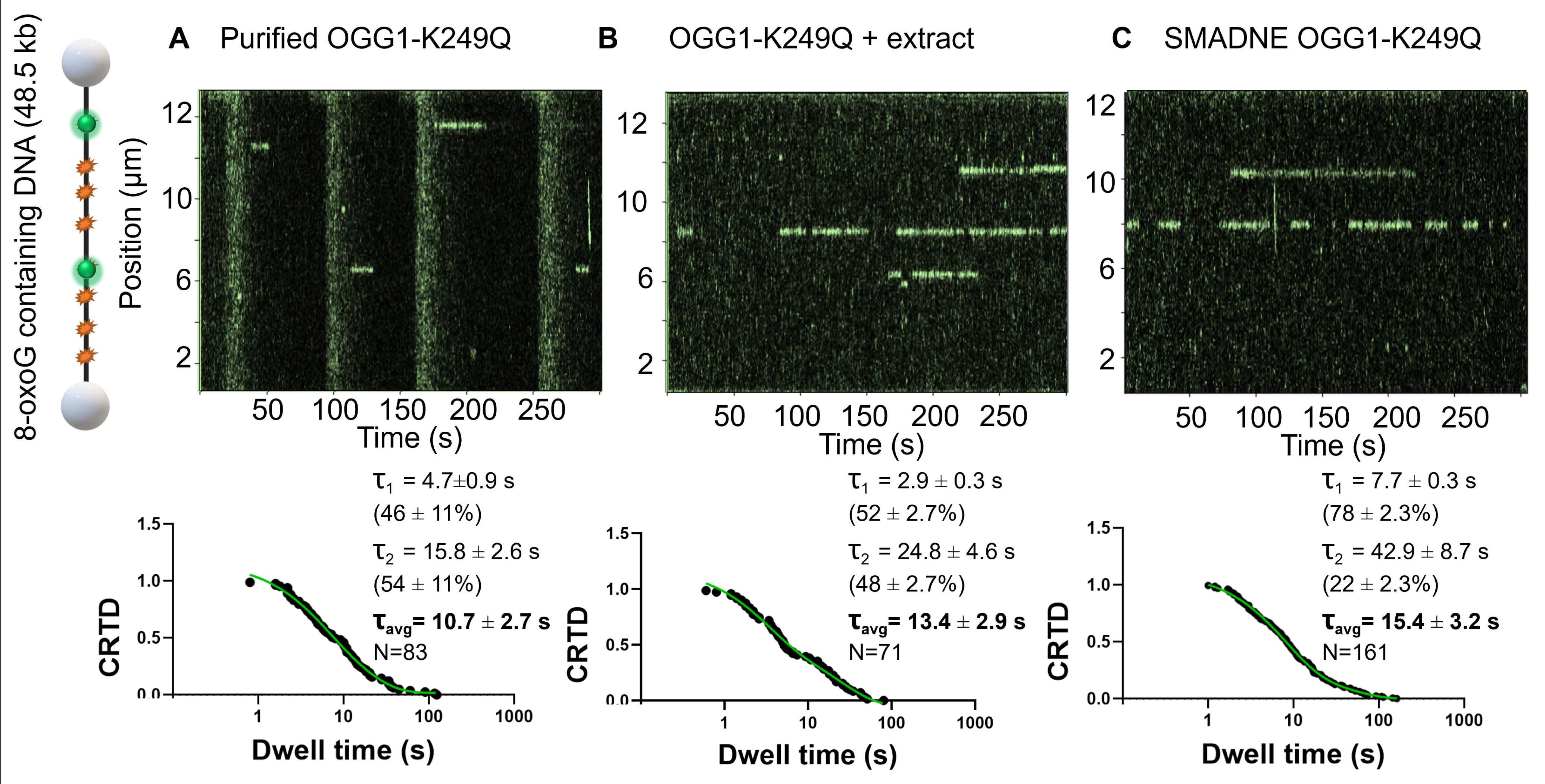
OGG1-K249Q binds 8-oxoG longer than WT as purified protein or with extract present. (A) Kymograph of OGG1-K249Q-GFP shown, with cartoon of streptavidin beads and DNA position shown on the left. The CRTD plot determined from the dwell times is shown beneath the kymograph. (B) OGG1-K249Q-GFP (green kymograph) also engages damage sites when in the presence of nuclear extracts. CRTD plot is displayed below. (C) Representative binding events from OGG1-K249Q-GFP events are shown, with the corresponding CRTD plot below.

## Discussion

Because the SMADNE approach [5] does not require protein purification, it promises to provide wider access to the single molecule regime. Nevertheless, it is essential to understand how the “dark” proteins in the extract may influence protein binding to DNA. The behavior of OGG1 was used as a test case and allowed for a direct comparison of: (1) OGG1-GFP purified from a bacterial expression system, (2) purified OGG1-GFP spiked into the nuclear extracts with no overexpressed protein, and (3) OGG1-GFP overexpressed using the SMADNE approach. The two latter conditions provide additional insight into the effects of dilute nuclear proteins (∼0.1 mg/mL) on the DNA binding behavior of a DNA binding protein. There are several considerations to keep in mind when selecting the best single-molecule approach, but selecting the best approach ultimately depends on the biological or biophysical question that is being addressed. Previous and ongoing single-molecule studies utilize purified proteins to yield robust results, reviewed in [11]. The SMADNE method allows a rapid characterization of WT and variant proteins and PTMs [5], increases binding specificity by reducing nonspecific binding, and allows biologically relevant facilitated dissociation to cause efficient release of proteins from their substrates (Fig. 5). In addition, mass spectrometry of the nuclear extracts indicated they contain chaperones that help stabilize proteins of interest, which will prove especially helpful for proteins that have disordered regions or low stability [5].

**Fig 5:**
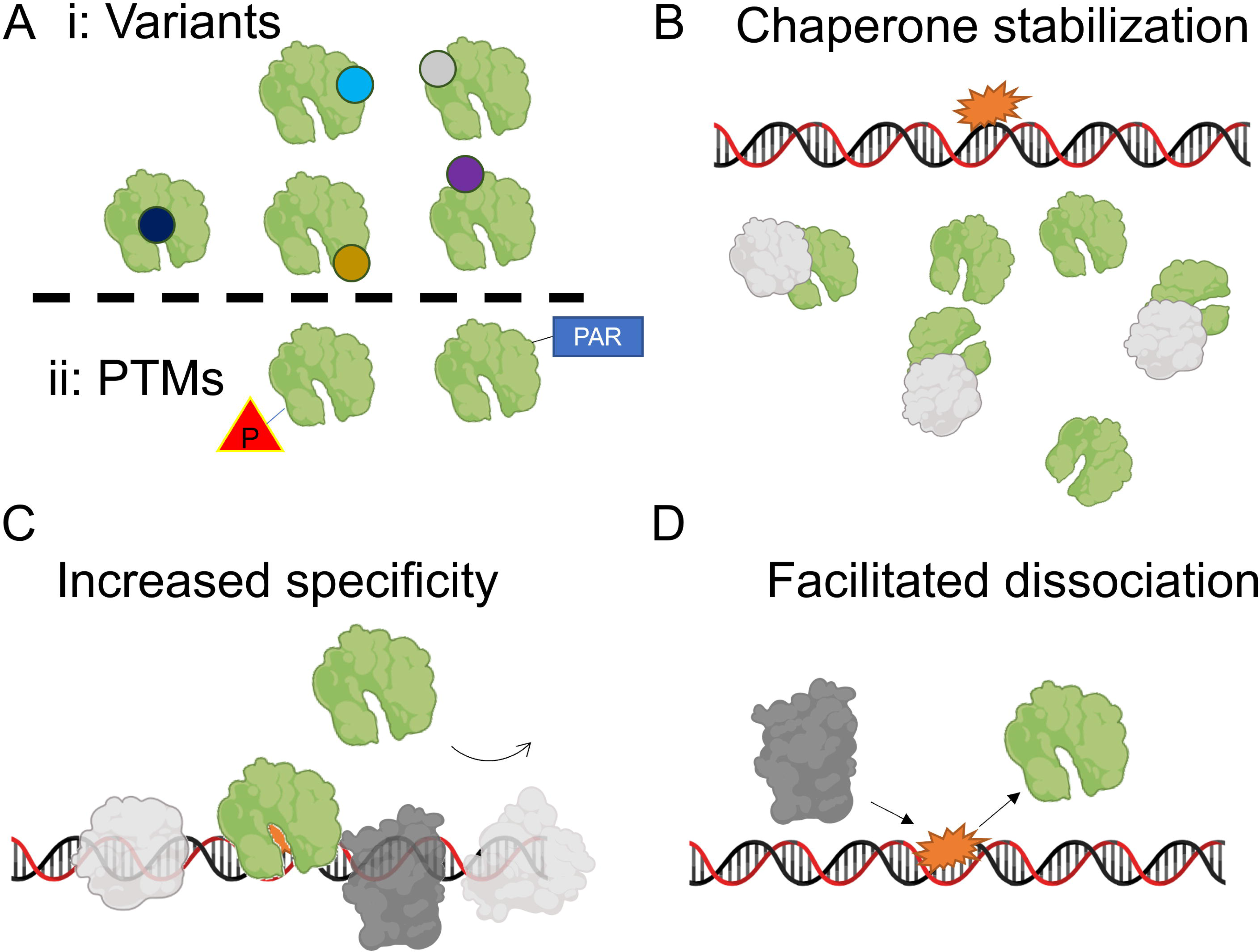
The roles of proteins in nuclear extracts on single-molecule analysis. (A) Nuclear extract approaches allow for variants (colored circles) and PTMs to be rapidly characterized. (B) Nuclear proteins (gray) increase data collection efficiency by stabilizing sample proteins (green) with chaperones and providing consistent functional protein concentrations. (C) Low-affinity engagement of nuclear proteins on undamaged DNA competes for nonspecific interactions of target proteins, increasing binding specificity. (D) Nuclear extract proteins assist in protein turnover on damage sites through a facilitated dissociation mechanism.

Because the SMADNE workflow is rapid (from plasmid to single-molecule data collection within three days), it allows the ability to quickly analyze variant proteins at the single-molecule level (Fig. 5A). These variants could be rationally designed to better understand the protein function, as in this present work, or even chosen from online databases to better understand how variants found in a clinical context contribute to function and thus disease. Many genes present in the Catalog of Somatic Mutations in Cancer (COSMIC, https://cancer.sanger.ac.uk/cosmic) have thousands of variants reported. SMADNE eliminates the necessity of protein purification and fluorescent labeling, democratizing single-molecule biophysical studies for a broad scientific community [25].

### Nuclear proteins in the SMADNE improve protein stability

Aside from workflow considerations, the other nuclear proteins present in the experimental conditions also offer other key advantages. In this study, we found that the concentrations of bacterially purified OGG1-GFP decreased over time, which causes difficulties in collection and analysis. This phenomenon may not occur with other purified protein and tag combinations, but the utilization of SMADNE can overcome this challenge should it arise. Secondly, chaperone proteins present in the nuclear extracts may increase the stability of proteins in the nuclear extract. Proteomic analysis of nuclear extracts made using the approach described here, indicated that two out of the top 20 most abundant proteins in the extract were identified as heat shock proteins (Heat shock protein HSP 90-beta and Heat shock cognate 71 kDa protein, see Table 2, Fig. 5B). Furthermore, SDS analysis of the major protein species in the extracts show consistently high reproducibility using the Abcam kit [5]. The levels of these chaperone proteins are on par with highly abundant nuclear proteins involved in nuclear structure, such as actin or nuclear pore complex protein Nup160. Thus, these and other chaperones likely stabilize proteins in solution during data collection; we routinely find that nuclear extracts can be utilized for hours of collection without apparent loss of activity. Furthermore, chaperones increasing protein stability could also explain why we observed a ∼3 second increase in weighted average binding lifetime for OGG1-K249Q-GFP present in nuclear extracts vs the purified protein alone. This stabilization phenomenon may be of significant importance when studying protein variants that disrupt protein stability.

**Table 2:** The 20 most abundant proteins present in nuclear extracts. Proteins that assist with protein folding are highlighted. Adapted from mass spectrometry experiment in [5].

| Protein names | Gene names | Mol. weight [kDa] |
| --- | --- | --- |
| Actin | ACTG1;ACTB | 41.792 |
| Annexin A2;Putative annexin A2-like protein | ANXA2;ANXA2P2 | 38.604 |
| Vimentin | VIM | 53.651 |
| Plectin | PLEC | 531.78 |
| Nuclear pore complex protein Nup160 | NUP160 | 162.12 |
| Filamin-A | FLNA | 280.74 |
| Annexin A1 | ANXA1 | 38.714 |
| Neuroblast differentiation-associated protein AHNAK | AHNAK | 629.09 |
| Annexin A5 | ANXA5 | 35.936 |
| Myosin-9 | MYH9 | 226.53 |
| ATP synthase subunit alpha, mitochondrial | ATP5A1 | 59.75 |
| Putative elongation factor 1-alpha-like 3;Elongation factor 1-alpha 1 | EEF1A1P5;EEF1A1 | 50.184 |
| ADP/ATP translocase 2;ADP/ATP translocase 2, N-terminally processed | SLC25A5 | 32.852 |
| ATP synthase subunit beta, mitochondrial | ATP5B | 56.559 |
| Heat shock protein HSP 90-beta | HSP90AB1 | 83.263 |
| Heat shock cognate 71 kDa protein | HSPA8 | 70.897 |
| Glyceraldehyde-3-phosphate dehydrogenase | GAPDH | 36.053 |
| Moesin | MSN | 67.819 |
| Kinesin-like protein KIF20B | KIF20B | 210.63 |
| Ezrin | EZR | 69.412 |

### Nuclear proteins in extract compete for undamaged DNA binding

One of the most striking differences between the purified OGG1 and OGG1 with nuclear extracts present was its behavior on undamaged DNA: numerous binding events on undamaged DNA were observed with purified OGG1, including some motile events that seemed to scan along the DNA. However, when the nuclear extracts are present these “nonspecific” events do not occur. Thus, unknown and unlabeled “dark” DNA binding proteins in the nuclear extract must bind the undamaged DNA and interfere with OGG1 binding (Fig. 5C). While we have not formally measured dark protein binding in this study, this is an active area of investigation and is necessary to unravel the full capabilities of the SMADNE approach. One potentially influential class of dark proteins are histones present in the nuclear extracts (detected by mass spectrometry) that may outcompete OGG1 for nonspecific DNA binding. Interestingly, however, the dark proteins do not seem to interfere with the ability of OGG1 to engage damage present on the DNA. Biologically speaking, other proteins blocking OGG1 from binding undamaged DNA may increase its damage-binding specificity, and this finding raises a question about whether OGG1 utilizes 1D diffusion in the nucleus for damage detection (where these dark proteins are presumably at much higher concentrations).The reduction in nonspecific binding could also present some challenges to the SMADNE approach: unless a protein exhibits a significantly high affinity to undamaged DNA, dark proteins binding to the DNA may limit the feasibility of fully characterizing the search mechanism of a DNA repair protein. It should be noted that other DNA repair proteins we have studied show non-specific DNA binding in nuclear extracts so this effect of “dark proteins” on OGG1 non-specific DNA binding might be protein specific, but is an important consideration when studying your protein of interest by SMADNE. To this end, 1D diffusion has been observed with the SMADNE approach for several other DNA repair proteins, including 3-alkyladenine DNA glycosylase (AAG) [26], thymine DNA glycosylase (TDG) [27], xeroderma pigmentosum complementation group C protein (XPC), and a variant of damaged-DNA binding protein 2 (DDB2) [5]. In studies of AAG, both the fraction of events that diffused and the rate of diffusion largely agreed between the data collected with nuclear extracts and the quantum dot-conjugated purified protein, suggesting that dark proteins did not alter the search process of AAG to the same extent as with the present study of OGG1.

### Proteins present in nuclear extracts may contribute to efficient repair mechanisms via facilitated dissociation

With purified proteins, the off-rate is generally believed to be independent of protein concentration [28]. However, the presence of unlabeled competitors can cause the off rate to increase due to the concept of facilitated dissociation [29–31]. In this phenomenon, the unlabeled proteins compete for sites on the DNA where the target protein has partially dissociated, and thus shift the equilibrium towards dissociation of the target. One advantage of utilizing GFP-fusion proteins in this study rather than conjugating the samples to Qdots or adding dyes to them with maleimide or N-hydroxysuccinimide reactions is that the fusion proteins are quantitatively labeled (that is, there is one fluorophore per protein and 100% of the purified proteins are labeled). In the purified context, this minimizes the possibility that unlabeled OGG1 can remove labeled protein once it has engaged the DNA. With the nuclear extracts, we did not utilize an OGG1 knockout cell line, so some endogenous OGG1 is present. However, with the overexpression of our fusion protein using a CMV promoter, we obtain expression levels 30-50 times higher than the endogenous protein, which translates to 97-98% labeled protein [5]. We previously did not see an impact of the endogenous protein until it reached about 25% of the level of the fluorescently tagged protein [5]. In an alternate approach to knocking down endogenous proteins, we have also successfully observed binding events for DNA repair factors when fluorescent tags are knocked in at the endogenous genes of interest thus preserving the ratio of unlabeled partners to target proteins and ensuring 100% of the protein of interest is label, (unpublished observation).

In nuclear extracts, however, several other proteins present in the extract could be assisting in OGG1 dissociation. Our group has specifically observed this phenomenon with UV-damaged DNA binding protein (UV-DDB), which stimulates the release of multiple DNA glycosylases from abasic sites, including OGG1 [18, 32], AAG [26], MUTYH [33], and SMUG1 [34]. Furthermore, endogenous apurinic/apyrimidinic endonuclease 1 (APE1) was also detected in nuclear extracts, which also has been shown to contribute to the efficient turnover of OGG1 [35]. In the present study, we observed evidence that nuclear proteins shortened the binding lifetime on DNA damage. In the experiments with WT OGG1-GFP on DNA with 8-oxoG, both purified OGG1-GFP resided longer on the DNA damage compared to purified OGG1-GFP spiked into nuclear extracts and OGG1-GFP generated by SMADNE. Thus, facilitated dissociation could be the mechanism by which the lifetimes are being shortened (Fig. 5D).

Interestingly, the WT OGG1-GFP expressed in mammalian cells exhibited a ∼three-fold shorter lifetime than the purified protein, suggesting that other factors may also be altering the binding lifetime. One potential factor could be the post-translational modification state of OGG1 when expressed in mammalian cells vs bacterial cells. OGG1 is modified numerous ways, and could be phosphorylated on a serine residue by protein kinase C [36], PARylated by PARP1 [37], acetylated by p300 [38], or even O-GlcNAcylated [39, 40]. These modifications are likely not made to the purified protein when it’s added to the extract because all of the cofactors needed for modification (NAD, ATP, and others) are greatly diluted during the nuclear extraction. Our measurements of NAD and ATP in undiluted nuclear extracts were approximately in the high nanomolar to 1 uM range. One other potential possibility is that the OGG1 protein could be at a different oxidation state when made in extracts vs purified from bacteria. One recent study found that OGG1 contains a nitrogen-oxygen-sulfur redox switch, and that the nitrogen from K249 contributes the nitrogen to the bridge[41]. The K249Q variant cannot form this bridge, which may explain why the purified variant protein spiked into extract condition exhibited a more similar lifetime to the SMADNE experiment compared to the WT protein where the switch was active. However, we also note that 1 mM fresh DTT was used in all experimental conditions, which may be enough to reduce any redox bridges present. An exciting prospect of the SMADNE approach is the ability to alter protein modifications prior to generating the nuclear extracts in order to address these questions about protein modification.

## Conclusions

The nucleus of a cell is a complex environment, with thousands of factors that could potentially impact the function of a single protein. Removing a protein from the milieu of a nucleus unlocks many potential techniques that are unattainable without purification, including structural studies and countless enzymological experiments. However, removing other nuclear factors from a protein comes at a cost, because purification pulls a protein of interest out of its native biological context. In biology, no protein works in isolation, and growing literature on pathway interplay implies that unexpected or even unknown proteins may assist in functions that are lost by purification. Directly analyzing proteins expressed in nuclear extracts at the single-molecule level represents an intermediate approach, through which new information can be gained that complements traditional biophysical experiments with purified proteins and cellular experiments. We believe SMADNE will provide a new window of observation into the behavior of nucleic acid binding proteins heretofore only accessible by biophysicists trained in protein purification and protein labeling. Furthermore, SMADNE will provide an opportunity for cell biologists who routinely study fluorescently tagged proteins in cell experiments to work within the single molecule regime.

## Acknowledgements

We thank Dr. Anna Campalans (Université de Paris-Cité) for the mammalian overexpression OGG1 plasmids, and lab members of for helpful discussion. We also thank Dr. Priya Raja and Dr. Ashna Nagpal for measurements of NAD and ATP in nuclear extracts, and Drs. Wim Vermeulen, Hannes Lans, Arjan Theil, and Jurgen Marteijn for providing knock-in cells for SMADNE analysis.

