## Supplementary Figures for "Single-molecule analysis of purified proteins and nuclear extracts: insights from 8-oxoguanine glycosylase 1"

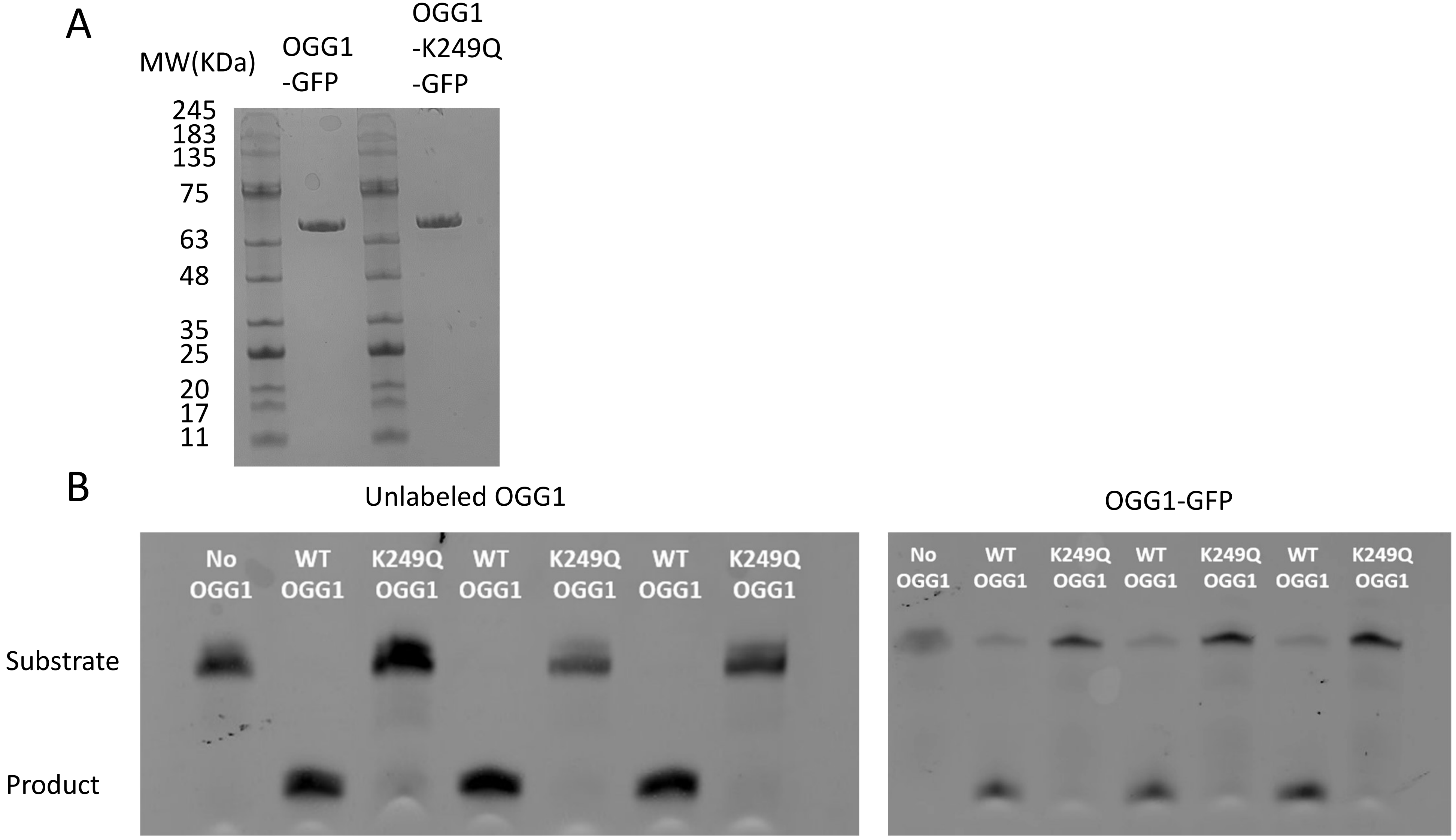
A
MW(KDa)
OGG1 -GFP
OGG1 -K249Q -GFP
245
183
135
75
63
48
35
25
20
17
11
B
Unlabeled OGG1
OGG1-GFP
No OGG1
WT OGG1
K249Q OGG1
WT OGG1
K249Q OGG1
WT OGG1
K249Q OGG1
Substrate
Product

### Fig S1 Legend

Fig. S1: OGG1-GFP purification and activity verification. (A) SDS-PAGE of purified OGG1-GFP and OGG1-K249Q-GFP, along with molecular standards labeled. (B) Enzymatic activity of OGG1-GFP (right) was compared to purified OGG1 without a label (left). With the GFP tag present, OGG1 still robustly cleaved 8-oxoG. 20 nM enzyme and 20 nM DNA substrate were incubated at 15 minutes in each lane.
